# AVM: a manually curated database of aerosol-transmitted virus mutations, human diseases, and drugs

**DOI:** 10.1101/2023.12.15.571955

**Authors:** Lan Mei, Yaopan Hou, Jiajun Zhou, Yetong Chang, Yuwei Liu, Di Wang, Yunpeng Zhang, Shangwei Ning, Xia Li

**Author notes:** Corresponding author(s). (Xia L), (Ning S), (Yun Z). Equal contribution (Lan Mei, Yaopan Hou).

## Abstract

Aerosol-transmitted viruses, with aerosol particles floating in the air to long distances, have the characteristics of strong infectivity and wide spread that is difficult to control. They cause various human diseases, posing a huge threat to human health. Some mutations can increase the transmissibility and virulence of the strain, which can reduce the protection of vaccines and weaken efficacy of antiviral drugs. Here, we established a manually curated database, AVM, to store this information. The current version of the AVM contains a total of 42,041 virus mutations, including 2613 immune escape mutations, 45 clinical information datasets, and 407 drugs, antibodies, or vaccines. In addition, we recorded 88 human diseases associated with viruses, and we found that the same virus can attack multiple target organs in the body and lead to diversified diseases. Further, the AVM database offers a straightforward user interface to expediently browse, retrieve, and download details. The AVM database is a comprehensive resource that provides timely and valuable assistance regarding the transmission, treatment, and related diseases of aerosol-transmitted viruses (http://bio-bigdata.hrbmu.edu.cn/AVM or http://www.bio-bigdata.center/AVM).

## Introduction

Aerosols are are collections of solid or liquid particles suspended in gases (such as air) [1], which may contain particles of any size. when droplets lose water in air suspension, proteins and pathogens composing the droplet nuclei are transmitted in aerosols. Aerosol-transmitted human viruses include a variety of viruses, such as severe acute respiratory syndrome coronavirus (SARS) [2], middle east respiratory syndrome coronavirus (MERS) [3], H1N1 [4], severe acute respiratory syndrome coronavirus 2 (SARS-2) [5], and Norovirus [6]. These viruses affect multiple organs, including the heart, kidney, liver, and small intestine, thereby causing serious diseases and great harm to human health. However, there is a shortage of integrated resources on aerosol-transmitted viruses.

Droplet nuclei (< 5 μm) can remain suspended in the air for an extended period of time, dispersing over long distances (> 1 m) [7]. Body secretions and excretions containing viruses can be atomized into droplets or particles containing infectious viruses by a variety of means, such as daily activities (e.g., talking, breathing, and coughing) [8], medical procedures (e.g., endotracheal intubation, noninvasive ventilation, and bronchoscopy) [9,10], and deposition of droplets on the surface of materials during human activities (walking, cleaning the room, and opening the door) [11]. Infectious aerosols are flooding our environment.

When the respiratory tract or body surface mucosa is exposed to infectious aerosols, the virus may destroy the host immune system and spread throughout the entire body, causing various diseases; however, the exact mechanism of viral transmission through aerosols is not fully understood. Some viral mutations (VMs) may have a significant effect on the development and progression of diseases by increasing or decreasing the pathogenicity or transmissibility of the virus [12,13]. For example, the mutation of P681H in *S* protein of COVID-19 leads to a structural alteration that enhances the affinity of furin binding to the *S* protein. This alteration facilitates the penetration of the virus into host cells, thereby augmenting its infectivity [14]. Mutants of the *S* gene with R685H substitution exhibit a significant decrease in the production of infectious virus, with 12- to 100-fold lower levels compared with those of wild-type (WT) virus. Additionally, the viral RNA levels of the *S* gene mutants are considerably lower than those observed in the WT virus [15]. Mutations in virus regions or genes may also increase the risk of disease progression and drug resistance [16,17]. A nonsynonymous change N74S in *NA* of H1N1 shows reduced susceptibility to zanamivir, thereby increasing the risk of drug resistance [16]. However, most of the data are scattered in many independent studies, and there is currently no systematic compilation of mutations of aerosol-transmitted viruses.

In the current public resources, much professional knowledge on viruses is stored in freely accessible databases. The Global Initiative on Sharing All Influenza Data (GISAID) [18], is the world’s largest influenza and novel coronavirus data platform for the submission of novel coronavirus and influenza virus genome sequence information. The Immune Epitope Database and Analysis Resource (IEDB) [19] documents the experimental data about antibodies and T-cell epitopes in human and animal studies. The Virus Pathogen Resource (ViPR) [20] is a comprehensive resource repository of data and analysis tools that cater to numerous virus families. The PATRIC [21] is dedicated to the development and integration of drug resistance gene database system. The GenBank [22] is a DNA-sequence database developed by the National Center for Biotechnology Information (NCBI). However, the existing databases lack compilation and integration of pathogenic VMs of aerosol-transmitted viruses and systematic evaluation and functional annotation of VMs. Such databases are especially needed because of increased concerns about various viruses that cause human disease in the wake of the COVID-19 outbreak.

Therefore, we constructed the AVM, a manually crafted database of human viruses transmitted via aerosols, which seeks to provide a comprehensive and professional resource of aerosol-transmitted viruses (**Figure 1**). The AVM involves mutation information, the effect of mutation on protein function, drug information, diseases, clinical data, and immune escape. This database offers experimentally validated data regarding the effect of VMs on protein function. It is the most comprehensive database on aerosol-transmitted viruses. The repository can be accessed at http://bio-bigdata.hrbmu.edu.cn/AVM or http://www.bio-bigdata.center/AVM.

**Figure 1.**
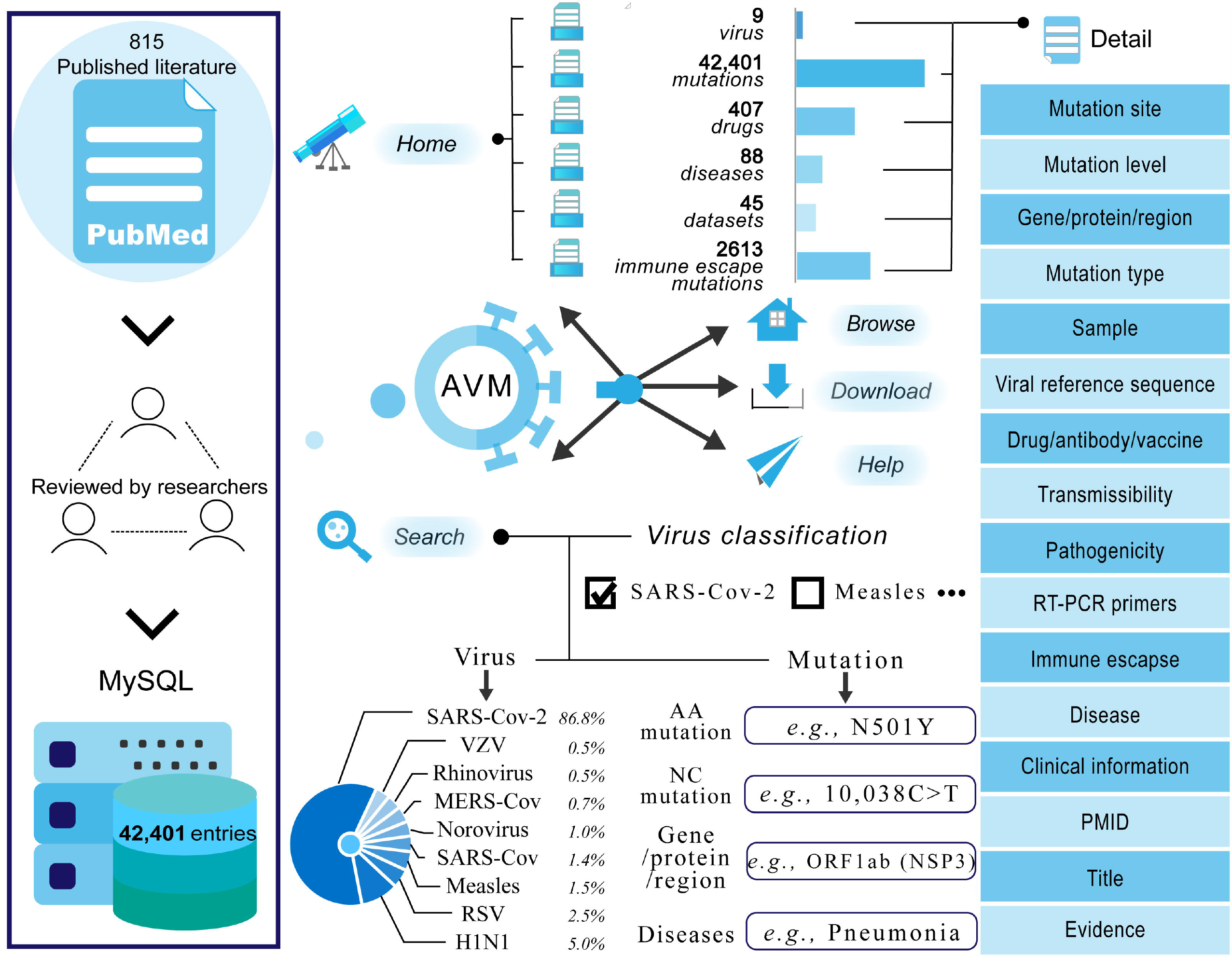
Data sources and the structure of AVM. SARS-Cov-2 (severe acute respiratory syndrome coronavirus 2), MERS-Cov (middle east respiratory syndrome coronavirus), SARS-Cov (severe acute respiratory syndrome coronavirus), Rhinovirus (human rhinovirus), VZV (Varicella-Zoster virus), RSV (respiratory syncytial virus), Measles (Measles virus), AA (amino acid), and NC (nucleotide).

### Evidence of virus transmission through aerosols

To assess the evidence of virus transmission through aerosols, we utilized the criteria outlined by Jones and Brosseau [23]. According to these criteria, aerosol transmission is considered plausible under the following conditions: (1) aerosols that contain the pathogen are produced by an infected individual; (2) the pathogen can remain active in the environment for a certain period; and (3) the aerosol has the potential to reach and infect target tissues. We assigned a value to each condition based on the strength of the evidence (1 for weak, 2 for moderate, and 3 for strong) and summed these values over the three conditions. Then, we deemed a score of 6 or higher to indicate the possibility of aerosol transmission. In the current version, the database incorporates a total of nine viruses, namely, MERS, RSV, Measles, Norovirus, Rhinovirus, VZV, SARS-2, SARS, and H1N1. They all adhere to the guideline proposed by Jones and Brosseau, and they have a score between 8 and 9 on a maximum of 9 (**Table 1**).

**Table 1.** Evidence of virus aerosol transmission.

| Virus | Aerosol generation | Viability in environment | Access to target tissue | Aerosol score | Ref. |
| --- | --- | --- | --- | --- | --- |
| RSV | A: Aerosols containing the pathogen are released from an infected person | A: The pathogen can linger in the air for a significant period of time while remaining surviving | B: Experimental infection by aerosol approach has been demonstrated in a mouse animal model | 8 | [24]<br>[25] |
| VZV | A: Aerosols containing the pathogen are emitted from an infectious person | A: In the previously occupied patient ward, the pathogen can be detected within 24 hours | A: The infection in humans by aerosol approach has been proven | 9 | [26]<br>[27] |
| MERS | A: Aerosols containing the pathogen are found in the air of isolation wards | A: The atomized virus has a long survival time in the environment, and the atomized virus is stable | B: Target tissues have been identified in human infection experiments through in vitro studies and non-aerosol routes, and transmission through aerosols seems plausible | 8 | [28]<br>[29]<br>[30] |
| SARS | A: Patients release aerosols containing the pathogen | A: Epidemiological evidence indicates transmission of aerosols over long distance | A: Experimental human infection via aerosol route has been confirmed | 9 | [31]<br>[32] |
| SARS-2 | A: Infected people release aerosols containing the pathogen | A: The pathogen remains infectious after surviving in aerosols around laboratory media for up to 16 hours | B: Experimental infection by aerosol approach has been demonstrated in hamster models | 8 | [33]<br>[34]<br>[35] |
| Norovirus | A: Air samples taken from the patient's room and toilet are positive, suggesting viral atomization in the patient's vomit | A: The pathogen can survive in the air for a certain period of time under suitable environmental conditions. Furthermore, in vitro aerosol studies have demonstrated that the pathogen retains the infectiousness and integrity | B: Target tissues have been identified in human infection experiments through non-aerosol approach. Alternatively, infection can be reasonably achieved through aerosol in vitro | 8 | [36]<br>[37]<br>[38] |
| H1N1 | A: The virus is emitted as an aerosol from infectious people | B: The pathogen around patients remains infectious after surviving in aerosols for several hours | A: Experimental human infection via aerosol route has been confirmed | 8 | [39]<br>[40]<br>[41] |
| Rhinovirus | A: Aerosols containing the pathogen are released from infected persons | A: Under specific air conditions, the pathogen using culture methods survives for several hours in laboratory media or fluids from the body | A: Experimental human infection via aerosol route has been verified | 9 | [42]<br>[43]<br>[44] |
| Measles | A: Aerosols containing the pathogen are released from infected persons | A: Under certain ambient conditions, the pathogen utilizing culture-based methods can remain viable for several hours in the air, whether in laboratory media or bodily fluids | B: Experimental infection by aerosol approach has been demonstrated in bovine animal models | 8 | [45]<br>[46]<br>[47] |

Note: A, B, and C represent the quality of evidence. A: strong, score: 3; B: moderate, score: 2; C: weak, score: 1. Aerosol score is the sum of the three evidence scores.

### Data collection and data processing

To maintain the quality of the AVM database in the collection process, each entry was meticulously gathered through a series of steps, which were employed for the construction of the VIMIC [48], ESC [49], and MSDD [50] databases.

At the outset, we searched the PubMed database [51] using the terms “*human* H1N1 mutation” and “*human* H1N1 variant.” The same approach was followed for all of the other viruses (e.g., *human* measles virus mutation and *human* measles virus variant; *human* norovirus mutation and *human* norovirus variant; and *human* respiratory syncytial virus mutation and *human* respiratory syncytial virus variant) to filter literature by matching keywords. We extracted key information by downloading all of the literature on VMs from peer-reviewed publications and preprints.

Second, we manually extracted information about VMs from academic articles. All of the chosen articles were evaluated by at least two researchers. During this stage, we recorded mutation site, mutation level (e.g., amino acid level and nucleotide level), mutation type (e.g., synonymous mutation and nonsynonymous mutation), virus gene, protein, and region, NCBI ID, virus genotype, subtype, and clade (genotypes or subtypes mentioned in the literature), drug, antibody, or vaccine (drug, antibody or vaccine mentioned in the literature), virus-associated disease, clinical information (clinical information available in the literature), virus sample (specimens derived from humans or cell lines), country (geographical distribution of the mutation), virus variants (virus subspecies), reference sequence, transmissibility (effect of mutations on viral transmissibility, e.g., promote or hinder), pathogenicity (effect of mutations on viral pathogenicity, e.g., increase or decrease), RT-PCR primers probes (yes or no), immune escape (mutation related to immune escape), evidence (e.g., a sentence containing information about the mutation in the literature), protein introduction (protein name, Uniprot ID, protein length, and protein description), and literature information (PubMed ID, year of publication, journal name, and paper title).

In addition, we searched the PubMed database for “H1N1 *human* disease” to obtain information on the human disease related to H1N1, and the same approach was followed for the other viruses (e.g., measles virus *human* disease, norovirus *human* disease, and respiratory syncytial virus *human* disease). The mode of transmissibility, pathogenicity, and immune escape caused by VMs were obtained from high-quality articles possessing robust experimental evidence. Finally, we used a standardized naming scheme, the Human Genome Variation Association rules (HGVS), to annotate each mutation.

### Database contents and construction

After completing this process, more than 810 articles were systematically reviewed, and a total of 42,041 entries, including 2613 immune escape mutations, 45 clinical information datasets, and 407 drugs, antibodies, or vaccines were manually curated (as of October 2, 2022). We also recorded 88 human diseases (**Table 2, Figure 2**). Moreover, we provided a brief description of each protein involved in mutation, including protein name, protein length, hyperlinks to Uniprot [52], and the structure and function of this protein. We also provided a brief description of the mechanism by which a mutation causes changes in transmissibility or pathogenicity of the virus from the original study.

**Table 2.**
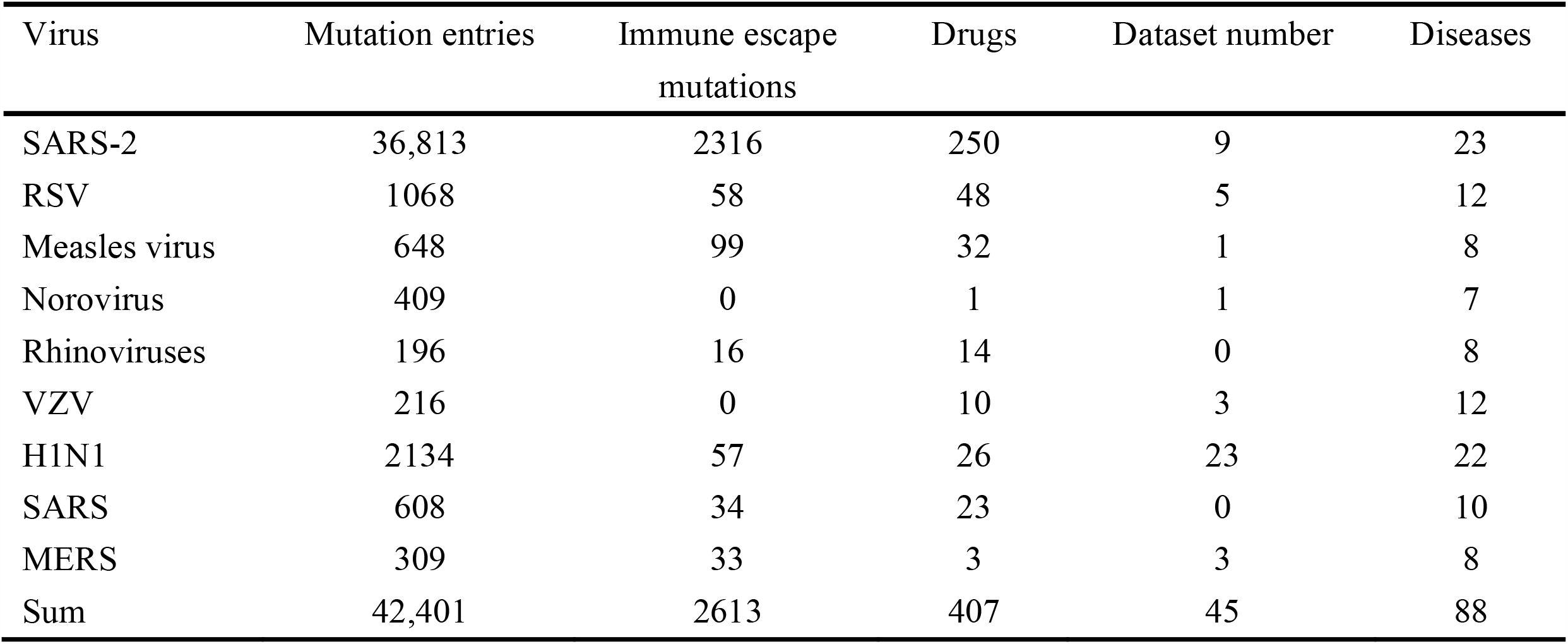
Statistics for the mutation entries in the AVM database.

| Virus | Mutation entries | Immune escape<br>mutations | Drugs | Dataset number | Diseases |
| --- | --- | --- | --- | --- | --- |
| SARS-2 | 36,813 | 2316 | 250 | 9 | 23 |
| RSV | 1068 | 58 | 48 | 5 | 12 |
| Measles virus | 648 | 99 | 32 | 1 | 8 |
| Norovirus | 409 | 0 | 1 | 1 | 7 |
| Rhinoviruses | 196 | 16 | 14 | 0 | 8 |
| VZV | 216 | 0 | 10 | 3 | 12 |
| H1N1 | 2134 | 57 | 26 | 23 | 22 |
| SARS | 608 | 34 | 23 | 0 | 10 |
| MERS | 309 | 33 | 3 | 3 | 8 |
| Sum | 42,401 | 2613 | 407 | 45 | 88 |

**Figure 2.**
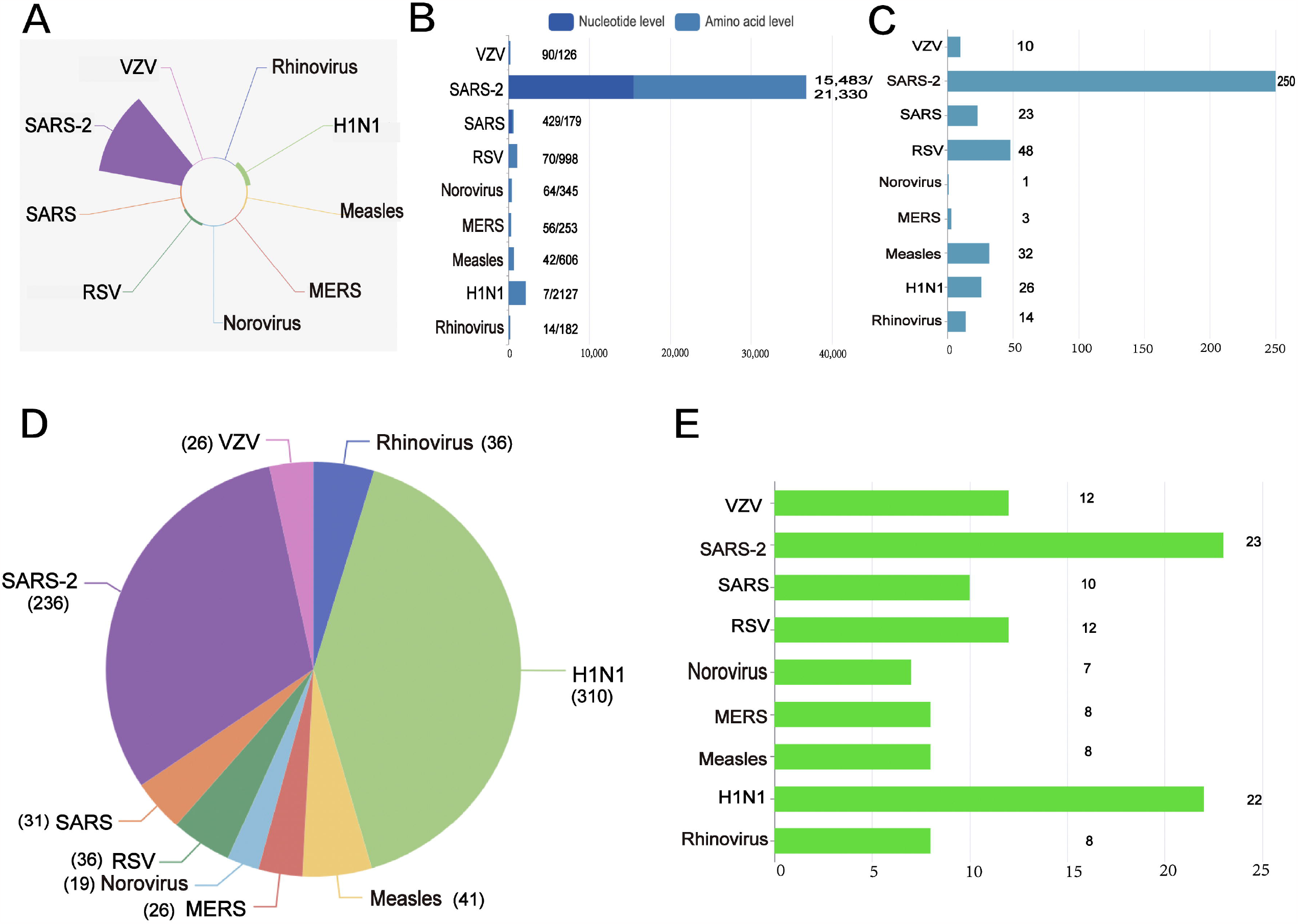
Information on AVM. **A**. The number of mutation entries for diverse virus. **B**. The number of NC mutations and AA mutations for diverse virus. The first number represents the count of NC level mutations, and the second number represents the count of AA level mutations. The two numbers are separated by “/” . **C**. The number of drugs, antibodies, or vaccines for diverse virus. **D**. The number of articles in PubMed for diverse virus. **E**. The number of human diseases caused by diverse viruses.

Ultimately, the entire data in the AVM were saved and administered by harnessing MySQL (version 8.0.17). JSP was utilized for constructing the web interfaces, and Java (version 1.8.0_291) was employed to code the data processing programs. The structure of web services was achieved using Apache Tomcat as the underlying framework. The AVM database can be accessed for free at http://bio-bigdata.hrbmu.edu.cn/AVM or http://www.bio-bigdata.center/AVM.

### User interface

The AVM renders an easy-to-use web interface that facilitates users to surf, search, and acquire data (**Figure 3**). The database presents a concise introduction to the virus genes and diseases caused by each virus on its respective interface, enabling users to swiftly verify their hypotheses about VMs.

**Figure 3.**
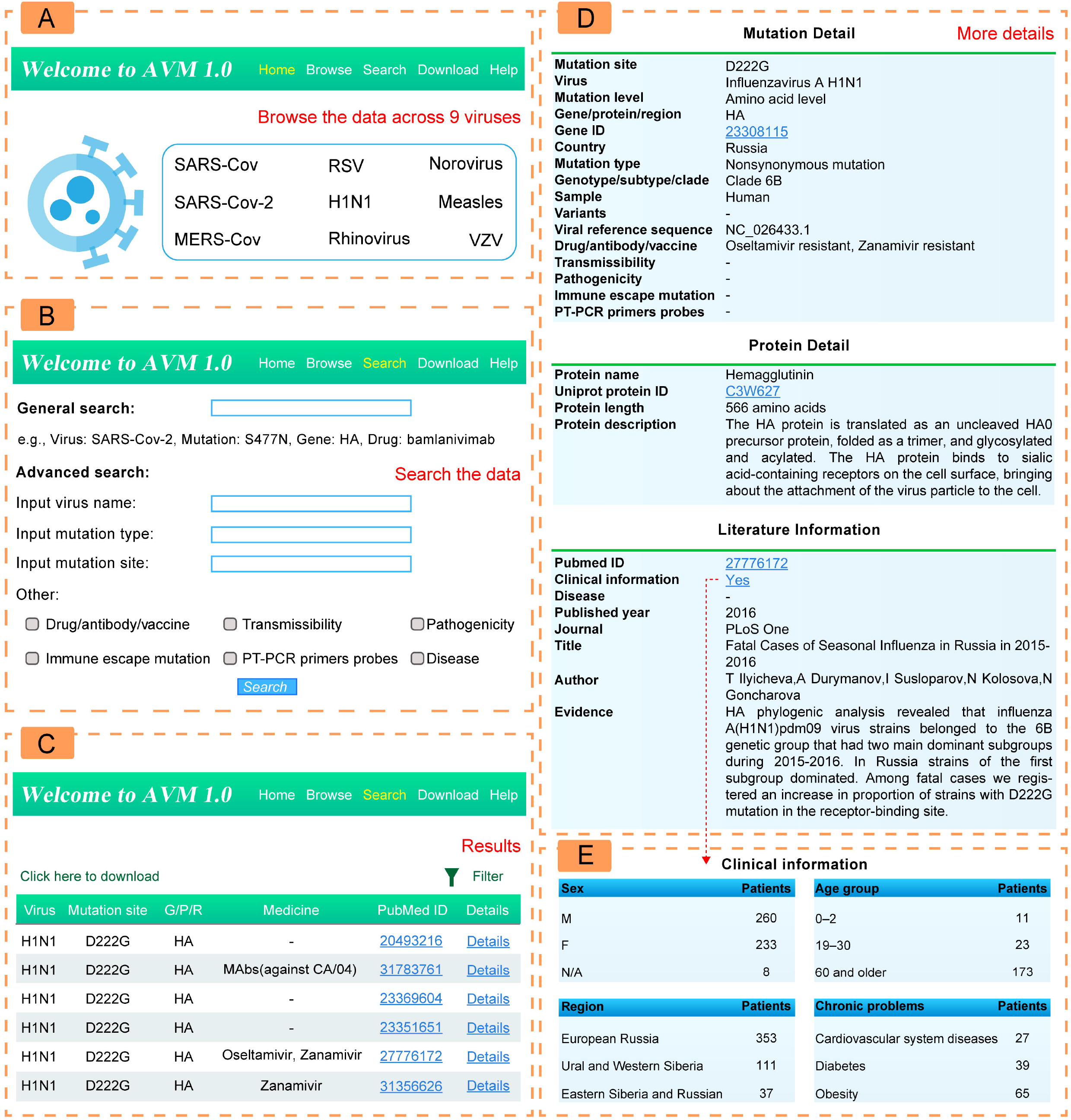
A schematic workflow of AVM. **A**. Users can click on some virus images to browse mutation information related to this virus. **B**. “Search” interface in a fuzzy and advanced manner. **C**. Results of the “Home” or “Search” page. **D**. Detailed information of the AVM entries. **E**. Clinical information of the AVM entries.

The statistics on the number of mutation entries for each virus are elaborated in the “Home” page. We also furnish a fuzzy search function. The users can click on a virus image and see relevant virus information; they can also search for a specific virus, VMs, some gene, and drugs. The users can consult a mutation site with mutation gene, protein, or region, drugs, and literature PMID. The detailed VMs page consists of mutation annotations, protein description, and literature information.

In the “Search” page, the AVM allows users to search by AA mutation, NC mutation, virus name, gene name, and drugs. The AVM also offers an advanced search, which allows users to screen for a drug, antibody, or vaccine mentioned in the literature, viral transmissibility, viral pathogenicity, RT-PCR probe binding sites, immune escape mutations, and diseases.

In the “Browse” page, we visually demonstrate the number of mutations of each virus in the database, the number of reference articles, the number of nucleotide and amino acid mutations, the number of related drugs and antibodies, and the diseases caused by the viruses. The users can download all of the data from the AVM on the “Download” page. The “Help” page provides a detailed tutorial explaining how the users can utilize the AVM effectively.

### Database application

#### Cases of transmissibility and pathogenicity

Owing to gathering a mass of experimentally demonstrated functional annotations of VMs associated with transmissibility and pathogenicity, the AVM could offer mechanistic notion and experimental basis for future research. As an illustration, searching the AVM for “A570D” (PMID: 34385690), a mutation in spike protein of SARS-2, we found that A570D hinders the spread of the virus. The description of transmission mechanism shows that A570D-mediated salt bridges could act as a pedal in a pedal-bin-like mechanism, controlling the motion of the *RBD*. This modulation is anticipated to affect *ACE2* binding and viral infectiousness. For another example, we searched the AVM for “L452R” (PMID: 34171266), in the *S* gene of SARS-2, which promotes virus transmission. The heightened viral infectivity resulting from the L452R substitution is strongly associated with an intensified electrostatic interaction with *ACE2*, likely because the residue 452 is situated near a cluster of negatively charged *ACE2* residues.

We also searched the AVM for “F49A” (PMID: 33999154), in *ORF1ab* (*NSP7*) of SARS-2, which decreases the pathogenicity of virus. The “Description Mechanism” displays that the mutations on the heterodimeric interface *I* of *NSP7* (*NSP7*−F49A and *NSP7*−L56A) and *NSP8* (*NSP8*−F92A) result in enhanced *NSP8* dimerization, leading to a decrease in the formation of the *NSP7*−*NSP8* heterotetramer. This indicates that these mutations likely disrupt the association between *NSP7* and *NSP8*. The AVM contains a large amount of mutation information about virus transmissibility and pathogenesis verified by experiments, which is beneficial for studying the virus transmission mechanism and pathogenicity and for further exploring the regulation mechanism of virus, thus making it possible to produce specific antiviral drugs.

#### Cases of immune escape and drug resistance

The AVM also provides the data of immune escape site and drug resistance site, along with the information on resistance antibody, vaccine, or resistance drug of the mutation. These data could contribute to and help professionals to choose a more scientific and reasonable treatment strategy, and also enable patients to achieve the best benefit through treatment. For example, by searching the AVM using “bamlanivimab resistant” in a general search, a commonly used monoclonal antibody drug for SARS-2 virus, we found that 32 entries such as A67V, D796Y, E484A, and E484K are resistant to bamlanivimab. Bamlanivimab should be avoided in clinical practice for patients with these mutations, and treatment strategies should consider dual antibody therapy or other monoclonal antibody therapy. As another example, we searched for “P653L” (PMID: 34061908), a mutation in *PA* of H1N1. The AVM shows that P653L is resistant to favipiravir. By searching the AVM for “favipiravir resistant” in a general search, another mutation of K229R in *PB1* appeared, which is also resistant to favipiravir. Further, the AVM will likely facilitate designing vaccines and neutralizing monoclonal antibodies, and guide the research and development of drugs. The AVM also includes a number of clinical datasets; hence, senior users could conveniently access the relevant virus data and conduct more in-depth analysis independently.

## Conclusion and future direction

In recent years, with the increasing popularity of sequencing technology, extensive virus biological data have been generated. A large number of VMs, especially in key genes or proteins of viruses, which lead to disease progression or drug resistance, have been found. Thus, we created the AVM, a database on aerosol transmitted viruses, which provides a comprehensive resource on VMs, drugs, and aerosol-transmitted human diseases. The AVM integrates experimental data on aerosol transmission of human pathogenic viruses, which is important to comprehend and even contain viral transmission.

The AVM will be further maintained and updated as the data of VMs accumulate in the future. We will also incorporate new analysis tools to consummate its practicability, add an upload module to empower users to submit VMs of interest, and augment RESTFUL API to keep pace of the research. In short, we hope that the AVM will help the biomedical community to obtain an all-encompassing comprehension of research related to viruses, and help to expedite the development of prophylactics and therapeutics for aerosol-transmitted viruses.

## Data availability

The AVM database can be accessed at http://bio-bigdata.hrbmu.edu.cn/AVM or http://www.bio-bigdata.center/AVM.

## CRediT author statement

**Lan Mei:** Methodology, Validation, Data curation, Visualization, Investigation, Writing - original draft. **Yaopan Hou:** Software, Visualization. **Jiajun Zhou:** Data curation, Investigation. **Yetong Chang:** Investigation. **Yuwei Liu:** Investigation. **Di Wang:** Investigation. **Yunpeng Zhang:** Supervision. **Shangwei Ning:** Supervision, Writing - review & editing, Funding acquisition. **Xia Li:** Conceptualization, Project administration. All authors have read and approved the final manuscript.

## Competing interests

The authors have declared no competing interests.

## Acknowledgments

This work received funding from the National Natural Science Foundation of China (grant numbers 62172131,32070673, 32070672), China Brain Project [2021ZD0202403]; Heilongjiang Touyan Innovation Team Program.; Outstanding Youth Foundation of Heilongjiang Province of China [YQ2021C026, YQ2022C034].

## References

[1] Leung NHL. Transmissibility and transmission of respiratory viruses. Nat Rev Microbiol 2021;19:528–45.

[2] Ding Y, He L, Zhang Q, Huang Z, Che X, Hou J, et al. Organ distribution of severe acute respiratory syndrome (SARS) associated coronavirus (SARS-CoV) in SARS patients: implications for pathogenesis and virus transmission pathways. J Pathol 2004;203:622–30.

[3] Ebrahim SH, Maher AD, Kanagasabai U, Alfaraj SH, Alzahrani NA, Alqahtani SA, et al. MERS-CoV Confirmation among 6,873 suspected persons and relevant Epidemiologic and Clinical Features, Saudi Arabia - 2014 to 2019. EClinicalMedicine 2021;41:101191.

[4] Syridou G, Drikos I, Vintila A, Pegkou A, Zografou L, Roungas P, et al. Influenza a H1N1 associated acute glomerulonephritis in an adolescent. IDCases 2020;19:e00659.

[5] Maffia-Bizzozero S, Cevallos C, Lenicov FR, Freiberger RN, Lopez CAM, Guano Toaquiza A, et al. Viable SARS-CoV-2 Omicron sub-variants isolated from autopsy tissues. Front Microbiol 2023;14:1192832.

[6] Li J, Wang B, He X, Li Z, Sun L, Li W, et al. Epidemiological characteristics of norovirus infection in pediatric patients during the COVID-19 pandemic. J Med Virol 2023;95:e28874.

[7] Ong SWX, Tan YK, Coleman KK, Tan BH, Leo YS, Wang DL, et al. Lack of viable severe acute respiratory coronavirus virus 2 (SARS-CoV-2) among PCR-positive air samples from hospital rooms and community isolation facilities. Infect Control Hosp Epidemiol 2021;42:1327–32.

[8] Alsved M, Nygren D, Thuresson S, Fraenkel CJ, Medstrand P, Löndahl J. Size distribution of exhaled aerosol particles containing SARS-CoV-2 RNA. Infect Dis (Lond) 2023;55:158–63.

[9] Tran K, Cimon K, Severn M, Pessoa-Silva CL, Conly J. Aerosol generating procedures and risk of transmission of acute respiratory infections to healthcare workers: a systematic review. PLoS One 2012;7:e35797.

[10] Zietsman M, Phan LT, Jones RM. Potential for occupational exposures to pathogens during bronchoscopy procedures. J Occup Environ Hyg 2019;16:707–16.

[11] Makison Booth C. Vomiting Larry: a simulated vomiting system for assessing environmental contamination from projectile vomiting related to norovirus infection. J Infect Prev 2014;15:176–80.

[12] Abduljaleel Z, Melebari S, Athar M, Dehlawi S, Udhaya Kumar S, Aziz SA, et al. SARS-CoV-2 vaccine breakthrough infections (VBI) by Omicron variant (B.1.1.529) and consequences in structural and functional impact. Cell Signal 2023;109:110798.

[13] Wang S, Xu X, Wei C, Li S, Zhao J, Zheng Y, et al. Molecular evolutionary characteristics of SARS-CoV-2 emerging in the United States. J Med Virol 2022;94:310–7.

[14] Mohammad A, Abubaker J, Al-Mulla F. Structural modelling of SARS-CoV-2 alpha variant (B.1.1.7) suggests enhanced furin binding and infectivity. Virus Res 2021;303:198522.

[15] Sasaki M, Toba S, Itakura Y, Chambaro HM, Kishimoto M, Tabata K, et al. SARS-CoV-2 bearing a mutation at the S1/S2 cleavage site exhibits attenuated virulence and confers protective immunity. mBio 2021;12:e0141521.

[16] Yates PJ, Mehta N, Horton J, Tisdale M. Virus susceptibility analyses from a phase IV clinical trial of inhaled zanamivir treatment in children infected with influenza. Antimicrob Agents Chemother 2013;57:1677–84.

[17] Zhang Y, Ndzouboukou JB, Lin X, Hou H, Wang F, Yuan L, et al. SARS-CoV-2 evolves to reduce but not abolish neutralizing action. J Med Virol 2023;95:e28207.

[18] Shu Y, McCauley J. GISAID: Global initiative on sharing all influenza data - from vision to reality. Euro Surveill 2017;22:30494.

[19] Vita R, Mahajan S, Overton JA, Dhanda SK, Martini S, Cantrell JR, et al. The Immune Epitope Database (IEDB): 2018 update. Nucleic Acids Res 2019;47:D339–43.

[20] Pickett BE, Sadat EL, Zhang Y, Noronha JM, Squires RB, Hunt V, et al. ViPR: an open bioinformatics database and analysis resource for virology research. Nucleic Acids Res 2012;40:D593–8.

[21] Antonopoulos DA, Assaf R, Aziz RK, Brettin T, Bun C, Conrad N, et al. PATRIC as a unique resource for studying antimicrobial resistance. Brief Bioinform 2019;20:1094–102.

[22] Benson DA, Cavanaugh M, Clark K, Karsch-Mizrachi I, Ostell J, Pruitt KD, et al. GenBank. Nucleic Acids Res 2018;46: D41–7.

[23] Jones RM, Brosseau LM. Aerosol transmission of infectious disease. J Occup Environ Med 2015;57:501–8.

[24] Aintablian N, Walpita P, Sawyer MH. Detection of Bordetella pertussis and respiratory synctial virus in air samples from hospital rooms. Infect Control Hosp Epidemiol 1998;19:918–23.

[25] Lindsley WG, Blachere FM, Davis KA, Pearce TA, Fisher MA, Khakoo R, et al. Distribution of airborne influenza virus and respiratory syncytial virus in an urgent care medical clinic. Clin Infect Dis 2010;50:693–8.

[26] Sawyer MH, Chamberlin CJ, Wu YN, Aintablian N, Wallace MR. Detection of varicella-zoster virus DNA in air samples from hospital rooms. J Infect Dis 1994;169:91–4.

[27] Gustafson TL, Lavely GB, Brawner ER, Jr., Hutcheson RH, Jr., Wright PF, Schaffner W. An outbreak of airborne nosocomial varicella. Pediatrics 1982;70:550–6.

[28] Kim SH, Chang SY, Sung M, Park JH, Bin Kim H, Lee H, et al. Extensive viable middle east respiratory syndrome (MERS) coronavirus contamination in Air and Surrounding Environment in MERS isolation wards. Clin Infect Dis 2016;63:363–9.

[29] van Doremalen N, Bushmaker T, Munster VJ. Stability of Middle East respiratory syndrome coronavirus (MERS-CoV) under different environmental conditions. Euro Surveill 2013;18:20590.

[30] Ki M. 2015 MERS outbreak in Korea: hospital-to-hospital transmission. Epidemiol Health 2015;37:e2015033.

[31] Tsai YH, Wan GH, Wu YK, Tsao KC. Airborne severe acute respiratory syndrome coronavirus concentrations in a negative-pressure isolation room. Infect Control Hosp Epidemiol 2006;27:523–5.

[32] Yu IT, Li Y, Wong TW, Tam W, Chan AT, Lee JH, et al. Evidence of airborne transmission of the severe acute respiratory syndrome virus. N Engl J Med 2004;350:1731–9.

[33] Liu Y, Ning Z, Chen Y, Guo M, Liu Y, Gali NK, et al. Aerodynamic analysis of SARS-CoV-2 in two Wuhan hospitals. Nature 2020;582:557–60.

[34] Fears AC, Klimstra WB, Duprex P, Hartman A, Weaver SC, Plante KS, et al. Persistence of severe acute respiratory syndrome coronavirus 2 in aerosol suspensions. Emerg Infect Dis 2020;26:2168–71.

[35] Sia SF, Yan LM, Chin AWH, Fung K, Choy KT, Wong AYL, et al. Pathogenesis and transmission of SARS-CoV-2 in golden hamsters. Nature 2020;583:834–8.

[36] Alsved M, Fraenkel CJ, Bohgard M, Widell A, Söderlund-Strand A, Lanbeck P, et al. Sources of airborne norovirus in hospital outbreaks. Clin Infect Dis 2020;70:2023–8.

[37] Bonifait L, Charlebois R, Vimont A, Turgeon N, Veillette M, Longtin Y, et al. Detection and quantification of airborne norovirus during outbreaks in healthcare facilities. Clin Infect Dis 2015;61:299–304.

[38] Atmar RL, Opekun AR, Gilger MA, Estes MK, Crawford SE, Neill FH, et al. Determination of the 50% human infectious dose for Norwalk virus. J Infect Dis 2014;209:1016–22.

[39] Lindsley WG, Noti JD, Blachere FM, Thewlis RE, Martin SB, Othumpangat S, et al. Viable influenza A virus in airborne particles from human coughs. J Occup Environ Hyg 2015;12:107–13.

[40] Bischoff WE, Swett K, Leng I, Peters TR. Exposure to influenza virus aerosols during routine patient care. J Infect Dis 2013;207:1037–46.

[41] Hao XY, Li FD, Lv Q, Xu YF, Han YL, Gao H. Establishment of BALB/C mouse models of influenza A H1N1 aerosol inhalation. J Med Virol 2019;91:1918–29.

[42] Huynh KN, Oliver BG, Stelzer S, Rawlinson WD, Tovey ER. A new method for sampling and detection of exhaled respiratory virus aerosols. Clin Infect Dis 2008;46:93–5.

[43] Karim YG, Ijaz MK, Sattar SA, Johnson-Lussenburg CM. Effect of relative humidity on the airborne survival of rhinovirus-14. Can J Microbiol 1985;31:1058–61.

[44] Couch RB, Cate TR, Douglas RG, Jr., Gerone PJ, Knight V. Effect of route of inoculation on experimental respiratory viral disease in volunteers and evidence for airborne transmission. Bacteriol Rev 1966;30:517–29.

[45] Bloch AB, Orenstein WA, Ewing WM, Spain WH, Mallison GF, Herrmann KL, et al. Measles outbreak in a pediatric practice: airborne transmission in an office setting. Pediatrics 1985;75:676–83.

[46] Ehresmann KR, Hedberg CW, Grimm MB, Norton CA, MacDonald KL, Osterholm MT. An outbreak of measles at an international sporting event with airborne transmission in a domed stadium. J Infect Dis 1995;171:679–83.

[47] Lemon K, de Vries RD, Mesman AW, McQuaid S, van Amerongen G, Yüksel S, et al. Early target cells of measles virus after aerosol infection of non-human primates. PLoS Pathog 2011;7:e1001263.

[48] Wang Y, Tong Y, Zhang Z, Zheng R, Huang D, Yang J, et al. ViMIC: a database of human disease-related virus mutations, integration sites and cis-effects. Nucleic Acids Res 2022;50:D918–27.

[49] Rophina M, Pandhare K, Shamnath A, Imran M, Jolly B, Scaria V. ESC: a comprehensive resource for SARS-CoV-2 immune escape variants. Nucleic Acids Res 2022;50:D771–6.

[50] Yue M, Zhou D, Zhi H, Wang P, Zhang Y, Gao Y, et al. MSDD: a manually curated database of experimentally supported associations among miRNAs, SNPs and human diseases. Nucleic Acids Res 2018;46:D181–5.

[51] Sayers EW, Bolton EE, Brister JR, Canese K, Chan J, Comeau DC, et al. Database resources of the national center for biotechnology information. Nucleic Acids Res 2022;50:D20–6.

[52] UniProt Consortium T. UniProt: the universal protein knowledgebase. Nucleic Acids Res 2018;46:2699.

